# Automated generation of personalized trajectories of aging phenotypes with DyViA-GAN

**DOI:** 10.1101/2025.08.22.671831

**Authors:** Saumyadipta Pyne, Deep Ray, Meghana S. Ray

## Abstract

With a general increase in human lifespan, the need for technological advances to develop strategies for healthy aging has assumed great importance. In the present study, our goal is to predict the progression of selected aging phenotypes in a given healthy individual as one continues aging past 65 years. Therefore, we developed a novel framework called Dynamic Views of Aging with conditional Generative Adversarial Networks (or DyViA-GAN) which is capable of predicting the plausible personalized trajectories of a selected aging phenotype conditioned on the available measurements of the phenotype at a few initial time instances, and additional covariates. Given the prevalence of osteoporosis in the aging population, we selected femoral neck Bone Mineral Density (BMD) of a healthy individual as the phenotype of interest, and baseline individual Body Mass Index (BMI) as covariate. We trained DyViA-GAN on a publicly available longitudinal dataset of a large cohort of mostly white women in the United States of age 65 years or above. Thus, it generated, for each individual, continuous phenotype trajectories, along with a corresponding region of acceptable predictions, for an age range of 66 to 89 years, for eight different combinations both with and without involving the covariate. The prediction results were subjected to rigorous quality-control and multiple comparative analyses. Our results clearly demonstrate the potential of generative deep learning frameworks in healthspan research.

## 1 Introduction

Forecasting the trajectories of diseases has been a cornerstone of the field of epidemiology involving numerous approaches ranging from the classical Susceptible-Infectious-Removed (SIR) and related models to more recent ones based on time series and predictive analytics, genomics, etc. [42]. In fact, modeling and prediction of infectious diseases have been among the key drivers of methodological advances in biostatistics and public health over the past century [44, 45]. Since the COVID-19 pandemic, novel approaches have emerged for outbreak forecasting including artificial intelligence (AI)-driven deep learning of spatio-temporal patterns, prediction by real and synthetic data fusion, e.g., [57, 18]. However, with a general increase in human lifespan over the past decades, some researchers in the field of disease prediction have now shifted their attention from the modeling of infectious disease outbreaks in a population to that of different chronic conditions and their progression in individuals in the form of personalized trajectories. It has led to such advances as deep aging clocks and various means of biological age assessment, e.g., [52, 16, 27, 48, 43, 41].

In the present study, our goal is to predict the progression of selected aging phenotypes in a given healthy individual as one continues aging past 65 years. Initial measurements of such phenotypes may be taken annually. With data from these initial time points as input, in combination with her other static or dynamic risk factors as covariates, we aim to build a reliable generative AI model that is capable of forecasting future values of the same phenotypes for this individual. Thus, our model will produce a set of plausible trajectories (univariate or multivariate depending on the selection of phenotypes), which could provide timely insights to inform and aid clinical decision-making that might extend her *healthspan*. For example, trajectories of aging phenotypes associated with one’s cognition or bone health can help one strategically build preparedness or adopt preventive measures against any future onset of dementia or possible fractures.

In particular, we focus on osteoporosis, which accounts for approximately 1.5 million fractures just in the U.S. each year [10]. According to its National Institute of Aging (NIA)*, osteoporosis is a “silent disease” since it does not have any markers of progression until a bone breaks – usually in the hip, spine. or wrist. In severe osteoporosis cases, a simple movement such as a cough or minor bump can result in a broken bone, i.e., a fracture. Additionally, individuals with osteoporosis may also have a longer duration of recovery and may experience chronic pain. In older adults, hip and spine fractures, in particular, can have serious consequences such as loss of mobility and independence. In a single typical year (2019), there were 318,797 emergency department visits, 290,130 hospitalizations, and 7,731 deaths related to hip fractures alone among adults of age 65 years or older in the U.S. [37]. Thus, osteoporosis acts as a key determinant of one’s (bone) healthspan over the course of aging.

Osteoporosis is defined as a systemic skeletal disease that has both the following characteristics: low bone mass and microarchitectural deterioration of bone tissues [39]. In clinical practice, osteoporosis is generally diagnosed when the measure of bone mineral density (BMD) of an individual is below the mean BMD of a reference population by at least 2.5 times the standard deviation, i.e., T-score ≤ −2.5 [30]. Low BMD or osteopenia is determined by a T-score between −1 and −2.5. As individuals age, BMD decreases and, consequently, osteoporosis becomes more prevalent in older adults [21, 46, 31]. A study of BMD of the femoral neck or lumbar spine estimated 43.4 and 10.2 million U.S. adults of age 50 years and above as having low bone mass and osteoporosis respectively.[55] While several factors may be involved in the etiology of osteoporosis, historically, the body mass index (BMI) has been linked to bone health as a protective factor [2, 46]. Paganini-Hill et al. using data from 8600 postmenopausal women reported that high BMI was associated with a significant reduction in hip fracture risk independently of other potential confounders [40]. Their findings were later corroborated by other studies [11, 14].

The longest running cohort study in the U.S. on this topic is the National Institute of Aging (NIA) sponsored Study of Osteoporotic Fractures (SOF), a multi-site study that ran from 1986 to 2017 and recruited over 10,000 participants at 4 different sites. Initially the study enrolled white women, and from 1997, it began to recruit African American women. Exclusion criteria included bilateral hip prostheses or inability to walk without assistance. Study participants attended a series of clinical visits with physical and mental evaluations including imaging to measure bone density, physical examinations, and questionnaires. In addition to clinical visits, participants were contacted via telephone or mail every four months to complete follow-up assessments for information on any falls, fractures, and vital status [1].

In a seminal paper in 1995, the SOF researchers established the dual energy X-ray absorptiometry (DEXA) scan as a non-invasive, painless, safe, and accurate measurement of BMD [11]. Subsequently, the U.S. Preventive Services Task Force osteoporosis management guidelines recommended routine BMD screening DEXA scans for women 65 years of age or older [38, 29]. The SOF study also showed that women who fell and broke their hip were five times more likely to die in their first year of fracture than those who did not break their hip [1]. Notably, beyond screening, DEXA scans can also serve to monitor osteoporosis progression, leading to opportunities for intervention and prevention of fractures. Therefore, femoral neck BMD data, as measured by DEXA, is an ideal initial choice of an aging phenotype to build a model for automated generation of personalized trajectories using a suitable computational framework.

Deep learning frameworks, such as Long Short-Term Memory Networks (LSTMs), Gated Recurrent Units (GRUs) and transformers networks, have been successful in applications involving time series forecasting [20, 50, 58]. The training and forecasting accuracy of these models typically usually rely on access to training data recorded over many time-points. Many of the previous studies that applied deep learning for producing disease trajectories used one or more recurrent neural networks (RNN) for predicting the ICD code for an individual’s next visit to the clinic [8, 9, 47]. Some have used generative adversarial networks (GANs) to predict such codes for multiple subsequent visits but not for tracking any actual phenotype of an individual [49]. Other deep learning applications have focused on specific health conditions or diseases, e.g., cystic fibrosis [32], Alzheimer’s disease [17]. Measures of DNA methylation biomarkers that estimate epigenetic aging serve a somewhat distinct purpose [24, 4]. Yet other platforms such as the Danish Disease Trajectory Browser (DTB) use statistical analysis of population-scale medical data to produce disease trajectories but these are not personalized predictions [51]. Various applications of generative AI and deep neural networks in aging research are described in reviews [54, 59] and the references therein.

To address such issues, we work with (a) individual longitudinal measurements of the selected phenotype that are available (b) only at a few (typically, 3-4) time-points. The aforementioned deep learning models are not appropriate to predict the phenotype trajectories under such conditions. We instead appeal to deep generative models which are suitable for solving probabilistic problems. Among existing generative frameworks, conditional Generative Adversarial Networks (cGANs) are popular in learning the underlying probability distribution of data conditioned on known values of other associated variables or parameters [35]. cGANs have been successfully used in studies of medical images such as for modeling the dynamics of cardiac aging [6], and Alzheimer’s disease progression based on MRI scans [26].

In this study, we proposed a novel cGAN framework called Dynamic Views of Aging with GAN (or DyViA-GAN) which is capable of predicting the plausible personalized trajectories of a selected aging phenotype conditioned on the available measurements of the phenotype at a few initial time instances, and additional covariates. For this purpose, we selected the BMD of the femoral neck of a healthy individual as the phenotype of interest and the baseline individual BMI as the covariate. Thus, DyViA-GAN generated personalized trajectories, for eight different combinations both with and without involving the covariate. The publicly available dataset used for training our model, due to the SOF study, is described in the Dataset section. Then we provided the details of the cGAN framework including the network architectures, and its model nomenclature for different data and covariate combinations in the Methods section. The predictions by DyViA-GAN were subjected to rigorous quality-control and multiple comparative analyses. The Results illustrate the corresponding cGAN-based generation of personalized trajectories of the selected phenotype. We end with Discussion on some of the limitations of our study and the scope for future work.

## 2 Methods

### 2.1 Dataset

We worked with an anonymized dataset for 9,704 mostly Caucasian women participants in the now concluded Study of Osteoporotic Fractures (SOF) study. The dataset is publicly available from the ‘SOF Online’ website (https://sofonline.ucsf.edu/). Attendance during the study varied as not all participant were available for all the visits, leading to significant gaps in the dataset. To ensure maximal number of participants in the training set, we considered participant data from Visit 2 (1989-90), Visit 4 (1992-94), Visit 5 (1995-96) and Visit 8 (2002-04), and considered the femoral neck BMD (denoted by FND) as measured by DEXA scans using Hologic QDR 1000 workstations. In accordance with best practices we exclude observations for patients over the age of 90 years to preserve patient anonymity [36]. We used longitudinally-adjusted scan measurements obtained by re-analyzing the previous scans and adjusting the regions-of-interest or deleting bone segments to match later scans. The T-scores of BMD were based on the parameters of a healthy reference population recommended by the International Society for Clinical Densitometry (ISCD) [33, 53].

In total, we have 2,113 de-identified participant records which were stratified based on the individual BMI values measured during Visit 1 (1986-87). We considered three strata: underweight and healthy participants (UWH) with BMI *<* 25, overweight participants (OW) with 25 ≤ BMI *<* 30, and obese participants (OB) with BMI ≥ 30. Details of the samples and stratification can be found in Table 1. We also used the recorded age of the various participants during each of the four visits, with the ages rounded down to half years.

**Table 1:** Description of data (FND T-scores) used for training and testing. Stratification based on underweight and healthy participants (UWH), overweight participants (OW), and obese participants (OB).

| | Strata | Number of Participants | Min. FND | Max. FND | Mean FND (± SD) |
| --- | --- | --- | --- | --- | --- |
| Training | UWH | 800 | -4.63 | 4.94 | -2.02 (± 0.87) |
| | OW | 720 | -5.07 | 2.51 | -1.69 (± 0.87) |
| | OB | 360 | -3.97 | 1.70 | -1.35 (± 0.92) |
| Test | UWH | 98 | -4.28 | 1.58 | -2.04 (± 0.94) |
| | OW | 95 | -4.03 | 5.42 | -1.70 (± 1.10) |
| | OB | 40 | -3.09 | 1.42 | -1.44 (± 0.96) |

### 2.2 DyViA-GAN

The proposed DyViA-GAN to predict personalized plausible trajectories of the selected phenotype comprises two neural networks, namely the generator 𝒢 and the discriminator 𝒟. We describe the algorithm for the most general setting where the phenotype of interest is denoted by *ρ* and additional known covariates is denoted by *Q*. Here *ρ* and *Q* may be a combination of different data modalities, such anthropometry, images, lifestyle, cognitive functions, physical performance, and biomarkers. We assume access to a dataset from which we can extract the measured values of the desired phenotype(s) *ρ* at two sets of time-points, {*t*_*i*_}_*i*∈ℐ_ and {*t*_*j*_}_*j*∈𝒥_, where ℐ and 𝒥 are two disjoint (non-overlapping) time indexing sets. The corresponding phenotype values are denoted by {*ρ*_*i*_}_*i*∈ℐ_ and {*ρ*_*j*_}_*j*∈𝒥_, respectively. Furthermore, the time-point sets {*t*_*i*_}_*i*∈ℐ_ and {*t*_*j*_}_*j*∈𝒥_ may be different and be unstructured across various records in the dataset.

The generator 𝒢 is fed as input the vector *Y* = (*Q*, {*ρ*_*i*_}_*i*∈ℐ_, {*t*_*i*_}_*i*∈ℐ_), a new time-point *t*, and a random latent variable *Z* which is sampled from an uncorrelated standard Gaussian distribution of dimension *N*_*z*_. The output of the generator is a predicted value of the phenotype in question at the time-point *t*.

We use the notations *X* = {*ρ*_*j*_}_*j*∈𝒥_ and *T*_*X*_ = {*t*_*j*_}_*j*∈𝒥_ to denote the “true” paired data associated with a given vector *Y*. The corresponding predictions by the generator are denoted by 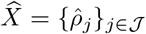 where 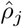 is the prediction at time-point *t*_*j*_. The scalar-valued discriminator network 𝒟 is fed as input either a “true” data tuple (*X, Y, T*_*X*_) or a “fake” tuple 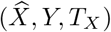. The role of the 𝒟 is to distinguish between true and fake tuples, while the role of the 𝒢 is to generate samples indistinguishable from those drawn from the true (unknown) conditional probability distribution of *X* given (*Y, T*_*X*_), i.e., 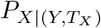.

The two network are trained in an adversarial fashion by solving the following optimization problem associated with a least-squares GAN [34]

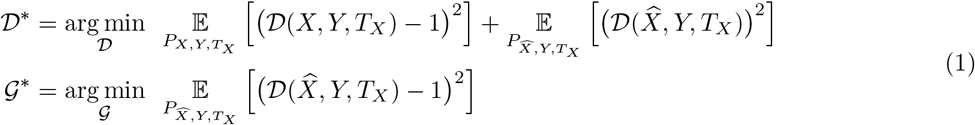

where 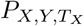 is the joint probability distribution of the true data, while 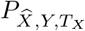 is joint probability distribution for the same data but with 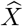 generated by 𝒢.

Once the DyViA-GAN is trained, the optimized generator 𝒢* is used to generate an ensemble of possible values of phenotype *ρ* at any point *t* given *Y* for an individual. This is achieved by drawing *K* random samples 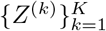 of the latent variable and evaluating 𝒢*(*Y, t, Z*^(*k*)^) for 1 ≤ *k* ≤ *K*. By allowing *t* to vary in the entire temporal window of interest, we can generate an ensemble of plausible continuous trajectories of phenotype for an individual. These trajectories can then be used to compute the point-wise empirical mean and variance of the predicted trajectories (given a *Y*) for any *t* as

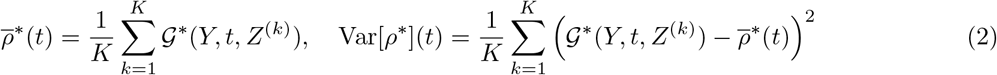

A schematic of the training and evaluation of DyViA-GAN is shown in Figure 1.

**Figure 1:**
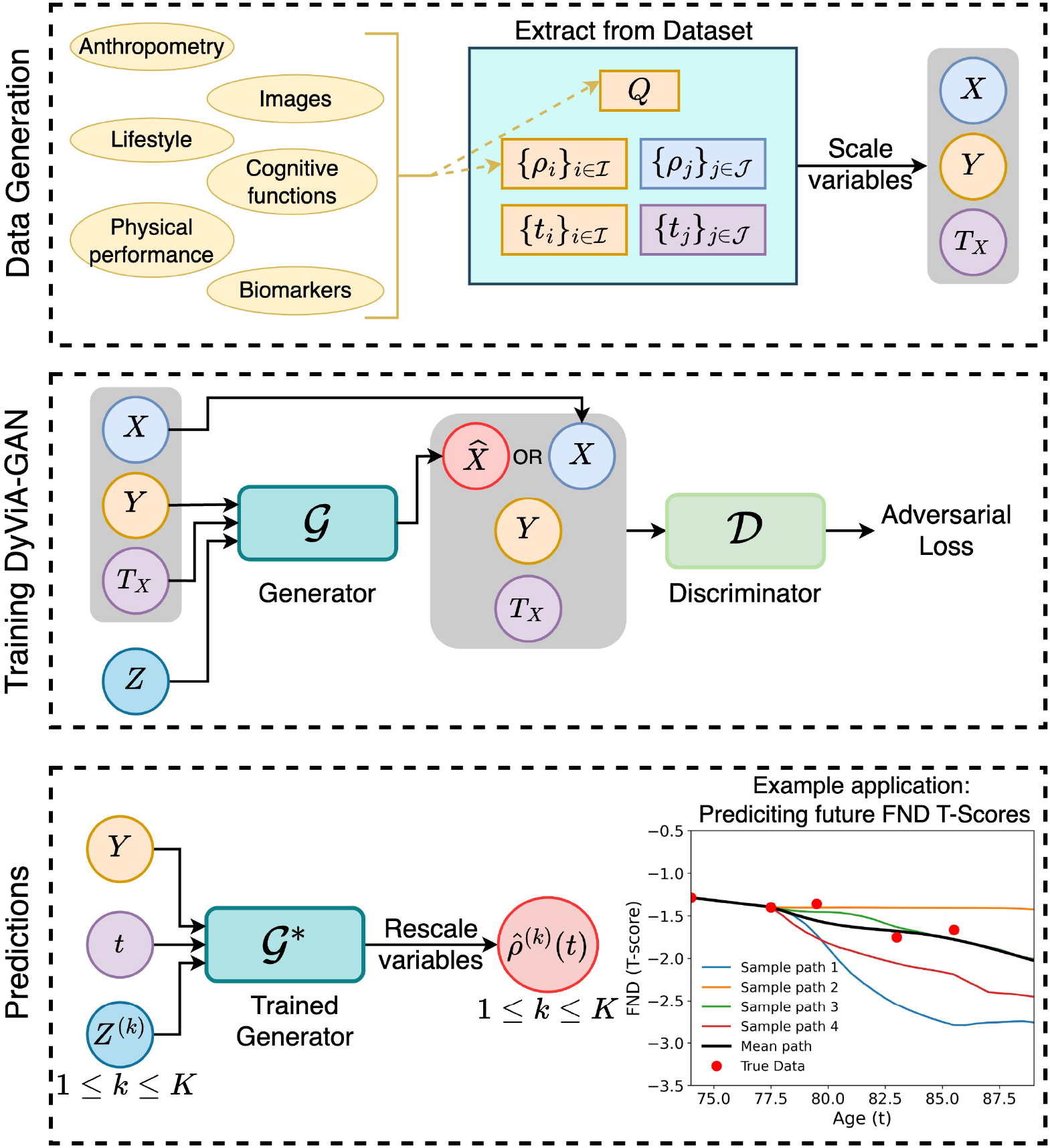
Schematic depicting the construction of the datasets, the inputs and outputs of 𝒢 and 𝒟 when training the DyViA-GAN, and how the trained generator 𝒢* is used to predict plausible phenotype trajectories. **Top Panel:** The phenotype measurements, the time-points of the measurements, and the additional covariates *Q* are extracted from the dataset to construct the tuple (*X, Y, T*_*X*_) after appropriate scaling. **Middle Panel:** During training phase, *Y, Z* and *T*_*X*_ are fed to 𝒢 to obtain the prediction vector 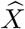 at the time-points in *T*_*X*_. The discriminator 𝒟 is fed either a true tuple (*X, Y, T*_*X*_) or a fake tuple 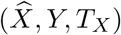. Both network are trained simultaneously according to Equation 1. **Bottom Panel:** The trained generator 𝒢* is fed the conditional data *Y* for a participant, any time-point *t*, and *K* samples of the latent variable. The output is an ensemble of *K* personalized trajectories 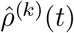 for 1 ≤ *k* ≤ *K* after rescaling the variables. As an example, we consider the FND prediction problem, with 4 sample trajectories and the mean trajectory (black curve) shown, which can be compared to the known phenotype measurements (red dots).

For the study on the SOF dataset considered in the present work, *ρ* is taken to be FND, while Q is chosen as the BMI measurement taken during Visit 1. Regression of the mean FND T-score against BMI in the data analyzed by this study shows a statistically significant positive association (*β* = 0.066, R^2^ = 0.116, p-value=0.017). Historically, it was shown that “A high body mass index protects against femoral neck osteoporosis in healthy elderly subjects [2].” As noted in subsection 2.1, we have FND values during Visits 2,3,5 and 8. Thus the index sets ℐ and 𝒥 used to construct *Y, X, T*_*X*_ will be chosen as subset of {2, 4, 5, 8} for training purposes. In particular, we set ℐ = {2, 4}, while 𝒥 = {5} or 𝒥 = {5, 8}. We scale the BMI by 100 before feeding it to the generator, and scale the age of the participants in accordance to

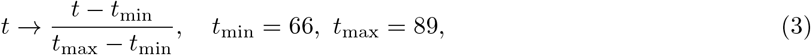

where [*t*_min_, *t*_max_] serves as the maximal age range (in years). These age limits were chosen based on the data available in the SOF dataset. The above scaling (3) ensures that *t* ∈ [0, 1] when fed as input to the network. We also remark here that the ages {*t*_2_, *t*_4_, *t*_5_, *t*_8_} differ across participants in the dataset, thus adding an additional layer of complexity to the trajectory prediction problem.

### 2.3 Network architectures

Let |ℐ| and |𝒥| denote the size of the index sets. For the choices made in the present work, |ℐ| = 2, while |𝒥| = 1 or 2. The generator 𝒢 is taken to be a fully-connected network with input dimension 2|ℐ| + *N*_*z*_ + 1 (or 2|ℐ| + *N*_*z*_ + 2 if BMI is used as a covariate) with a scalar-valued output. We used four hidden layers of width 50 each with the hat activation function [23, 56]. To to ensure that the generator’s prediction is constrained to pass through the given initial measurements corresponding to ℐ, i.e., at *t*_2_ and *t*_4_, the output of the generator is transformed as follows

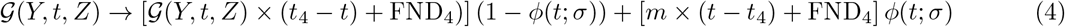

where

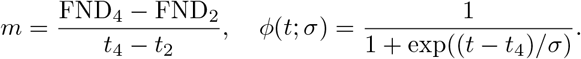

The function *ϕ*(*t*; *σ*) in the above transformation smoothly blends the linear curve passing through (*t*_2_, FND_2_) and (*t*_4_, FND_4_) to the network output. In this work, we choose *σ* = 10^−2^.

The discriminator 𝒟 is a also taken to be fully-connected network with input dimension 2|ℐ| + 2|𝒥 | (or 2|ℐ| + 2|𝒥 | + 1 if BMI is used as a covariate) with a scalar-valued output. We used four hidden layers of width 50 with the hat activation function. Other hyperparameter values for DyViA-GAN are listed in Table 2.

**Table 2:** Common hyperparameter values used for the DyViA-GAN models.

| Hyperparameter | Value/Choice |
| --- | --- |
| Latent Dimension $N_z$ | 20 |
| Optimizer | Adam |
| Learning rate for $\mathcal{G}$ | $10^{-3}$ |
| Learning rate for $\mathcal{D}$ | $10^{-3}$ |
| Number of $\mathcal{D}$ updates per $\mathcal{G}$ update | 4 |
| $\ell^2$ regularization parameter | $10^{-7}$ |
| Batch size | ALL: 200, OWH: 200, OW: 180, OB: 90 |
| Total epochs | 2000 |

### 2.4 DyViA-GAN model nomenclature

We trained multiple version of DyViA-GAN on different data and covariate combinations. For easier reference, we use the nomenclature Model-⟨DT⟩ for the models trained on different data types determined by tag ⟨DT⟩. The tag value ⟨DT⟩ = ALL implies that the model is trained on unstratified training data with 2100 samples (see Table 1), ⟨DT⟩ = UWH implies the networks are trained on the UWH strata samples with 900 samples, ⟨DT⟩ = OW implies training on the OW strata with 800 training samples, while ⟨DT⟩ = OB implies training on the OB strata with 400 training samples. Furthermore, the model name ends with an additional tag -Q if the covariate *Q* is fed as input to the generator. For example, Model-OWH denotes the model trained (and tested) on the OWH strata, while Model-OB-Q denotes the model trained (and tested) on OB strata with *Q* used as an input for the generator. In all, we consider 8 DyViA-GAN models: 1) Model-ALL, 2) Model-ALL-Q, 3) Model-UWH, 4) Model-UWH-Q, 5) Model-OW, 6) Model-OW-Q, 7) Model-OB, and 8) Model-OB-Q. Furthermore, we consider two variants of the model, one with 𝒥 = {5} and another with 𝒥 = {5, 8}. Thus there are 8 DyViA-GAN models for each variant.

### 2.5 Evaluation of model performance

Using the trained generator 𝒢*, we quantify the performance of the various models via the following metrics:

- **Root mean squared error (RMSE)**: This will be evaluated either across all time-points not fed as input to the generator (see Table 3), or at specific time-points (see Table 4).
- **Maximal score:** For a particular participant, we define the score 𝒮(*t*) at any time-point *t* using the empirical measures computed in (2)

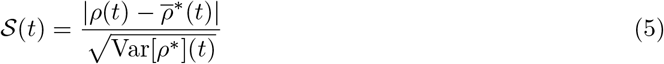

which is the normalized distance of the true value of the phenotype at *t* from the mean predicted value at that time-point. The maximal score 𝒮_*M*_ is define as the maximum over of an individuals score over all time-points not shown to the generator, and at which the true value is known. In the present context, 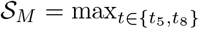 *S*(*t*), with the empirical statistics computed using *K* = 1000 random samples of *Z*. Thus, predictions with lower maximal scores are preferable. We say that the *score is acceptable* for a sample if 𝒮_*M*_ ≤ *ϵ*_*s*_ where *ϵ*_*s*_ is a tunable parameter.
- **Filtering plausible trajectories:** In order to eliminate outliers from the predicted plausible trajectories for an individual, we perform an additional post-processing filtering. For each predicted plausible trajectory 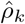, 1 ≤ *k* ≤ *K*, consider the mean path 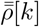 computed using the remaining *K* − 1 trajectories, i.e., excluding the *k*-th trajectory. We then retain the *k*-th trajectory if dist 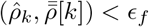, where dist() is a meaningful function to measure the distance between trajectories, while *ϵ*_*f*_ is a tunable parameter. In the present work, we take dist() to be the Dynamic Time Warping (DTW) distance, which is a commonly used elastic shape-based measure of similarity between a pair of time series [15].

**Table 3:** Percentage (%) of samples with acceptable scores (𝒮_*M*_ ≤ 2.5) for 8 different DyViA-GAN models using 𝒥 = {5} (top subtable) and 𝒥 = {5, 8} bottom subtable). The percentages are also listed for each data strata for models trained on unstratified samples. For models trained on stratified data, the percentages are listed only for the corresponding strata. The notation - indicates no scores are evaluated for a particular strata with a given model.

| Models using $\mathcal{J} = \{5\}$ | | | | | | | | |
| --- | --- | --- | --- | --- | --- | --- | --- | --- |
| Model \ Strata | Training |  |  |  | Test |  |  |  |
|  | UWH | OW | OB | ALL | UWH | OW | OB | ALL |
| Model-ALL | 89.12 | 86.81 | 88.61 | 88.14 | 87.76 | 83.16 | 90.00 | 86.27 |
| Model-ALL-Q | 89.75 | 86.94 | 88.61 | 88.46 | 87.76 | 84.21 | 90.00 | 86.70 |
| Model-UWH | 85.00 | - | - | - | 77.55 | - | - | - |
| Model-UWH-Q | 91.62 | - | - | - | 87.76 | - | - | - |
| Model-OW | - | 84.72 | - | - | - | 82.11 | - | - |
| Model-OW-Q | - | 90.56 | - | - | - | 90.53 | - | - |
| Model-OB | - | - | 79.17 | - | - | - | 80.00 | - |
| Model-OB-Q | - | - | 84.17 | - | - | - | 85.00 | - |

| Models using $\mathcal{J} = \{5, 8\}$ | | | | | | | | |
| --- | --- | --- | --- | --- | --- | --- | --- | --- |
| Model \ Strata | Training |  |  |  | Test |  |  |  |
|  | UWH | OW | OB | ALL | UWH | OW | OB | ALL |
| Model-ALL | 82.62 | 78.33 | 81.11 | 80.69 | 82.65 | 80.00 | 87.50 | 82.40 |
| Model-ALL-Q | 80.00 | 76.11 | 79.44 | 78.40 | 79.59 | 78.95 | 87.50 | 80.69 |
| Model-UWH | 79.88 | - | - | - | 77.55 | - | - | - |
| Model-UWH-Q | 81.50 | - | - | - | 79.59 | - | - | - |
| Model-OW | - | 72.78 | - | - | - | 72.63 | - | - |
| Model-OW-Q | - | 90.56 | - | - | - | 90.53 | - | - |
| Model-OB | - | - | 72.22 | - | - | - | 77.50 | - |
| Model-OB-Q | - | - | 73.33 | - | - | - | 80.00 | - |

**Table 4:** RMSE for 8 different DyViA-GAN models using 𝒥 = {5} (top subtable) and 𝒥 = {5, 8} bottom subtable). The errors are also listed for each data strata for models trained on unstratified samples. For models trained on stratified data, the errors are listed only for the corresponding strata. The notation - indicates no errors are evaluated for a particular strata with a given model.

| Models using $\mathcal{J} = \{5\}$ | | | | | | | | |
| --- | --- | --- | --- | --- | --- | --- | --- | --- |
| Strata<br>Model | Training |  |  |  | Test |  |  |  |
|  | UWH | OW | OB | ALL | UWH | OW | OB | ALL |
| Model-ALL | 0.53 | 0.57 | 0.52 | 0.54 | 0.56 | 0.68 | 0.57 | 0.61 |
| Model-ALL-Q | 0.55 | 0.59 | 0.53 | 0.56 | 0.59 | 0.69 | 0.58 | 0.63 |
| Model-UWH | 0.52 | - | - | - | 0.56 | - | - | - |
| Model-UWH-Q | 0.55 | - | - | - | 0.59 | - | - | - |
| Model-OW | - | 0.55 | - | - | - | 0.65 | - | - |
| Model-OW-Q | - | 0.55 | - | - | - | 0.66 | - | - |
| Model-OB | - | - | 0.59 | - | - | - | 0.64 | - |
| Model-OB-Q | - | - | 0.63 | - | - | - | 0.68 | - |

| Models using $\mathcal{J} = \{5, 8\}$ | | | | | | | | |
| --- | --- | --- | --- | --- | --- | --- | --- | --- |
| Strata<br>Model | Training |  |  |  | Test |  |  |  |
|  | UWH | OW | OB | ALL | UWH | OW | OB | ALL |
| Model-ALL | 0.44 | 0.50 | 0.47 | 0.47 | 0.48 | 0.61 | 0.51 | 0.54 |
| Model-ALL-Q | 0.38 | 0.44 | 0.43 | 0.41 | 0.43 | 0.56 | 0.46 | 0.49 |
| Model-UWH | 0.38 | - | - | - | 0.43 | - | - | - |
| Model-UWH-Q | 0.39 | - | - | - | 0.44 | - | - | - |
| Model-OW | - | 0.45 | - | - | - | 0.56 | - | - |
| Model-OW-Q | - | 0.51 | - | - | - | 0.60 | - | - |
| Model-OB | - | - | 0.46 | - | - | - | 0.50 | - |
| Model-OB-Q | - | - | 0.44 | - | - | - | 0.48 | - |

We remark that the parameters *ϵ*_*s*_ used to define acceptable scores and *ϵ*_*f*_ used filter trajectories are both are problem-dependent and tunable by the practitioner based on domain expertise. In the present work, we choose *ϵ*_*s*_ = 2.5 and *ϵ*_*f*_ = 2.0 motivated by the ablation study presented in Appendix B.

## 3 Results

We ran DyViA-GAN on the individual samples of available (or “true”) FND measurements from the described dataset to generate their respective personalized continuous trajectories of FND based on the stated models. Using the criteria discussed in (subsection 2.5), we evaluate and compare the performance of the various designed models. Although Model-ALL and Model-ALL-Q are trained on unstratified data, we also evaluate the trained generators on stratified data for a more fine-grained analysis. The RMSE and maximal scores of the other models trained on stratified data are only evaluated on the corresponding strata.

### 3.1 Score-based comparison

When comparing the two variants of the model, we observe from Table 3 that a larger percentage of training and test samples have acceptable scores with models trained using 𝒥 = {5}, as compared to models trained using 𝒥 = {5, 8}. This indicates that the plausible trajectories generated using the first variant have a larger spread about the predicted mean trajectory, thus having a better chance of capturing dynamics with larger temporal variations. We also observe that feeding the covariate Q (i.e., the BMI) as a generator input improves the performance of the models (both variants) trained on stratified data. For instance, the percentage of test samples with acceptable scores with models trained on OW strata increases from 82.11% to 90.53% for the first variant of the model (see top subtable in Table 3) and from 72.63% to 90.53% for the second variant of the model (see bottom subtable in Table 3).

### 3.2 RMSE-based comparison

If we consider the mean predicted trajectory as a single representation of the phenotype evolution, then the RMSE is a more appropriate metric to measure model performance. We observe from Table 4 that the models trained with 𝒥 = {5} (top subtable) consistently lead to higher RMSE when compared to the respective models trained with 𝒥 = {5, 8} (bottom subtable). Thus, based on the RMSE metric, the later variant of the DyViA-GAN models perform significantly better. In general, the inclusion of the BMI as a generator input seems to marginally deteriorate the RMSE – with the exception of the 𝒥 = {5, 8} variant of Model-ALL-Q where the error reduces when BMI is included.

### 3.3 Checking and filtering predicted trajectories

With the goal of eliminating outliers among the predicted trajectories, we filter the trajectories generated by the DyViA-GAN models using DTW with a filtering parameter *ϵ*_*f*_ = 2.0. We demonstrate the qualitative performance of the trained models with this filtering on two test participants chosen from each strata UWH, OW and OB. For each participant, we generate *K* = 1000 trajectories using each model, and then compute the personalized mean trajectory only using the filtered trajectories. The filtered trajectories and corresponding mean trajectories are shown in Figure 2 (OWH), Figure 3 (OW) and Figure 4 (OB).

**Figure 2:**
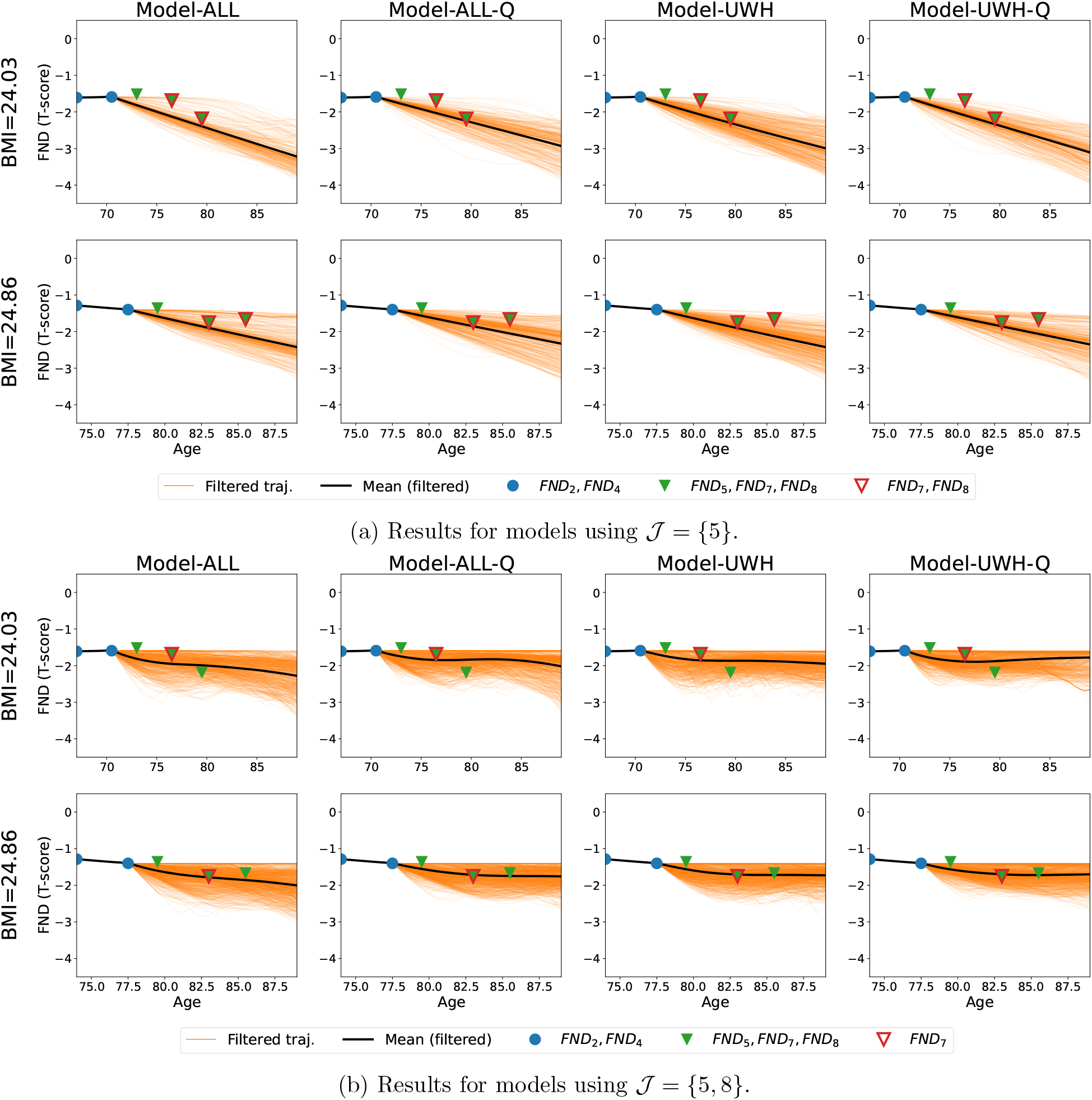
Predictions with various models for two test participants from strata UWH. Each plot depicts the filtered trajectories (orange curves) out of 1000 original trajectories, the mean filtered personalized phenotype trajectory (black curve), and the phenotype measurements at times *t*_2_, *t*_4_, *t*_5_, *t*_7_, *t*_8_ (dots and triangles). We depict the generator input using blue dots, the generator predictions using green triangles, and mark the prediction at Visit 7 (never shown to the DyViA-GAN) using red (unfilled) triangles. The BMI of the participant is listed on the left of each row.

**Figure 3:**
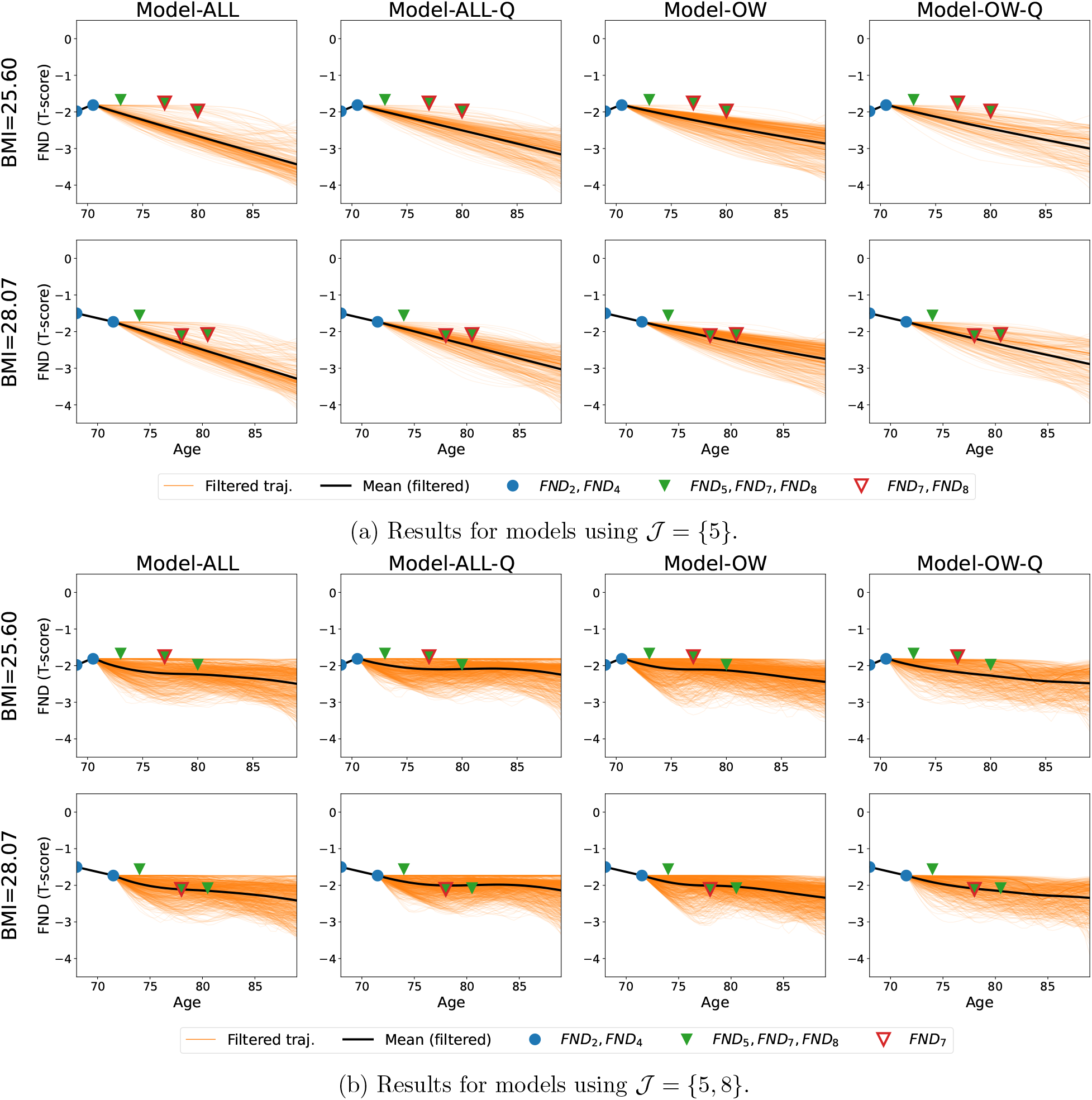
Predictions with various models for two test participants from strata OW. Each plot depicts the filtered trajectories (orange curves) out of 1000 original trajectories, the mean filtered personalized phenotype trajectory (black curve), and the phenotype measurements at times *t*_2_, *t*_4_, *t*_5_, *t*_7_, *t*_8_ (dots and triangles). We depict the generator input using blue dots, the generator predictions using green triangles, and mark the prediction at Visit 7 (never shown to the DyViA-GAN) using red (unfilled) triangles. The BMI of the participant is listed on the left of each row.

**Figure 4:**
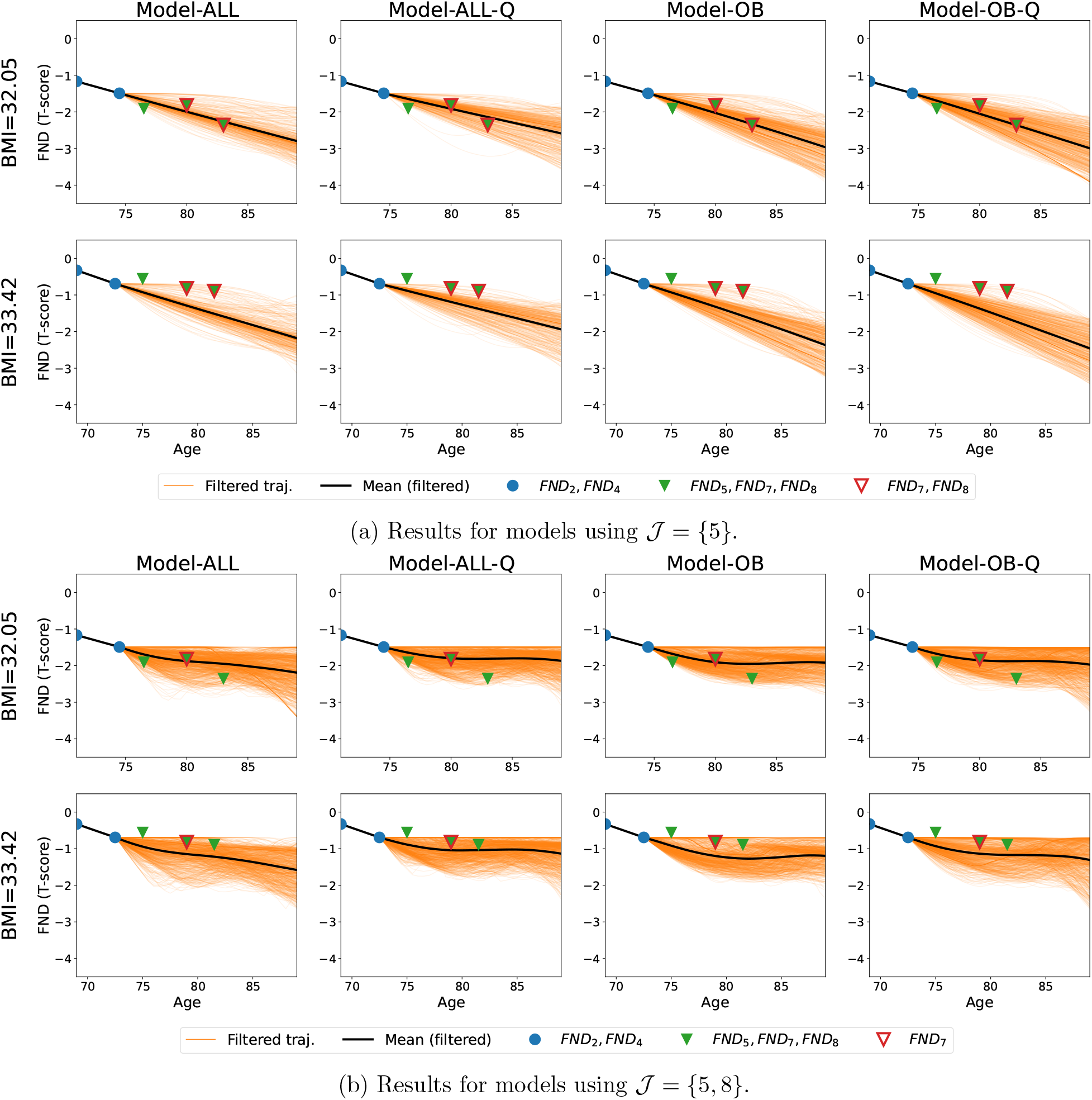
Predictions with various models for two test participants from strata OB. Each plot depicts the filtered trajectories (orange curves) out of 1000 original trajectories, the mean filtered personalized phenotype trajectory (black curve), and the phenotype measurements at times *t*_2_, *t*_4_, *t*_5_, *t*_7_, *t*_8_ (dots and triangles). We depict the generator input using blue dots, the generator predictions using green triangles, and mark the prediction at Visit 7 (never shown to the DyViA-GAN) using red (unfilled) triangles. The BMI of the participant is listed on the left of each row.

In addition, we have access to the adjusted phenotype value at Visit 7 (1999) for these six test participants. This serves as held-out data point for both variants of DyViA-GAN, as data from this visit was not used during the training phase. Note that Visit 8 is also a held-out data point for the models trained with 𝒥 = {5}. We depict the generator inputs using blue dots, the generator predictions using green triangles, and mark the unseen/held-out time-points using red (unfilled) triangles in Figure 2, Figure 3 and Figure 4. Appendix A contains additional plots depicted a few sample trajectories for each of these test participants for better clarity. We make the following observations from the plots in Figure 2, Figure 3 and Figure 4:

- The mean trajectories pass through the true data at *t*_2_, *t*_4_, which is a feature of the constraint Equation 4 applied on the generator output.
- With the filtering threshold *ϵ*_*f*_ = 2.0, a significantly larger number of plausible trajectories are accepted with the 𝒥 = {5, 8} model variants as compared to the 𝒥 = {5} variants. This can be easily inferred by the observing the denseness of the filtered trajectories shown in the figures.
- The filtered trajectories with the 𝒥 = {5, 8} models are significantly more nonlinear with richer structure, while those obtained using the 𝒥 = {5} models appear to be mostly (piecewise) linear (also see Supplementary Material).
- The (filtered) mean with the 𝒥 = {5, 8} models leads to better approximation of the data at unseen data at future time point *t*_7_.
- The mean trajectory with all models generally predict a downward trend in the FND values as time progresses, which is consistent with clinical observations in aging patients.

### 3.4 Benchmarking against other methods

We compare the proposed DyViA-GAN against other standard approaches. In particular, we consider a standard cubic smoothing spline model, linear mixed-effect models (LMMs), and Gaussian process (GP) regression. Among these, the later two approaches are well suited to for longitudinal data with unstructured time-points.

#### 3.4.1 Cubic splines

We use a standard cubic smoothing spline model [19] as implemented by the ‘splinefun’ function in R. It has been noted that the advantage of such a model (say, over a full ARIMA model) is that not only it provides a smooth historical trend but also a linear forecasting ability, which is hardly affected by the restricted parameter space [25]. The dataset to train the spline model is constructed by considering the maximal number of participants in the SOF dataset who have FND measurements from Visits 2, 4, 5 and 8 while also sharing the same ages *t*_2_, *t*_4_, *t*_5_ and *t*_8_ during these visits. This leads to 49 participants with (unscaled) ages 69.0, 72.5, 74.5 and 81.5 years. The spline model was trained on the first two ages and then use to predict the T-scores during the last two ages. We compare the RMSE of the spline predictions with the mean predictions at these two ages using the various DyViA-GAN models trained previously on unstratified data, i.e., Model-ALL and Model-ALL-Q. As show in Table 5, the DyViA-GAN models outperform the spline model when predicting the T-scores at 74.5 years. However, only the 𝒥 = {5, 8} variants lead to lower errors than splines at age 81.5 years. Furthermore, including BMI as input improves the performance of this latter variant, which is also consistent with the observations from Table 4.

**Table 5:**
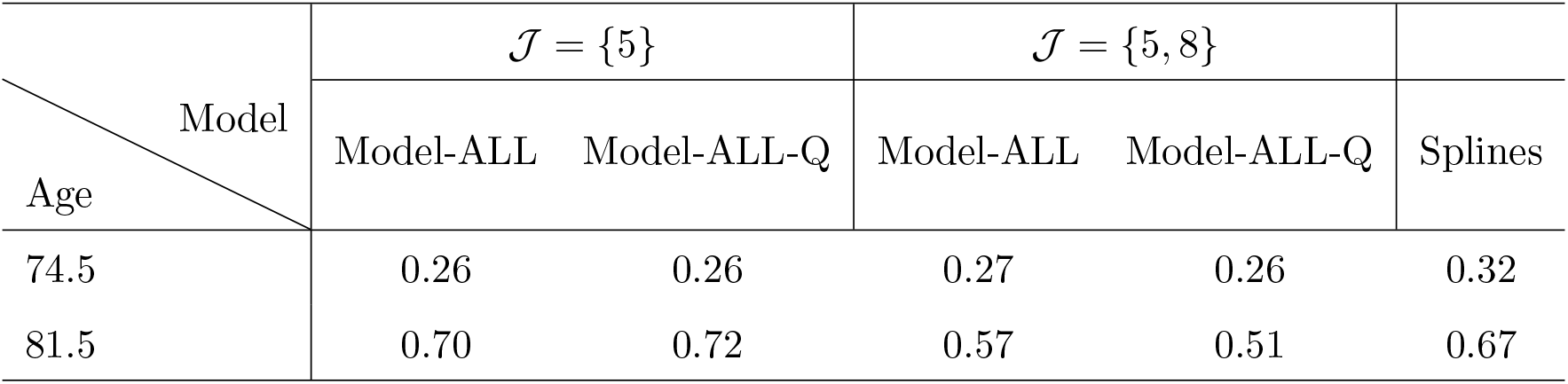
Comparing the performance of Model-ALL and Model-ALL-Q with a spline-based regression model. The RMSE of the predictions at ages 74.5 and 81.5 years are shown.

#### 3.4.2 Linear mixed-effects models

We consider LMMs with fixed (population-level) effects modeled using a polynomial basis (in time) of dimension *p* and random (individual-level) effects modeled using a polynomial basis of dimension *q*. The random basis coefficients are modeled using a normal distribution 𝒩(0, Σ), and additional (individual-level) observation noise is modeled using 𝒩(0, *σ*^2^*I*). The coefficients of the fixed basis, covariance Σ for the random coefficients, and *σ*^2^ for the noise model are determined by maximizing the log-likelihood on the (unstratified) training data. Once trained, the predictions for an individual are conditioned on their measurements at the first two time-points. We use the nomenclature LMM (p,q) to denote the LMM trained with basis dimension *p* and *q*.

We consider three variants of LMM based on the values of *p* and *q*. As shown in Table 6, the RMSE (of 0.49) on the test samples remains fairly robust across all variants, which perform better than Model-ALL and at par with Model-ALL-Q. We also compare the performance of the models on the six test participants considered in the previous sections. We observe from Figure 5 that all LMMs predict fairly linear (mean) trajectories. However, this is not ideal when the individual-level evolution of phenotypes is non-monotonic, as can be seen in Figure 5 for participants with BMI 25.60 and 32.06. In comparison, the DyViA-GAN predictions are far more expressive and capable of capturing non-monotone behavior. Furthermore, we have the flexibility of choosing a network architecture that ensures the predictions pass through (or close to) the given observations, which need not hold true for LMMs.

**Table 6:** RMSE of the predictions of various models on the unstratified test data.

| Model | RMSE |
| --- | --- |
| LMM (3,2) | 0.49 |
| LMM (6,2) | 0.49 |
| LMM (6,4) | 0.49 |
| GP Const. | 0.62 |
| GP Lin. | 0.61 |
| GP Quad. | 0.62 |
| Model-ALL | 0.54 |
| Model-ALL-Q | 0.49 |

**Figure 5:**
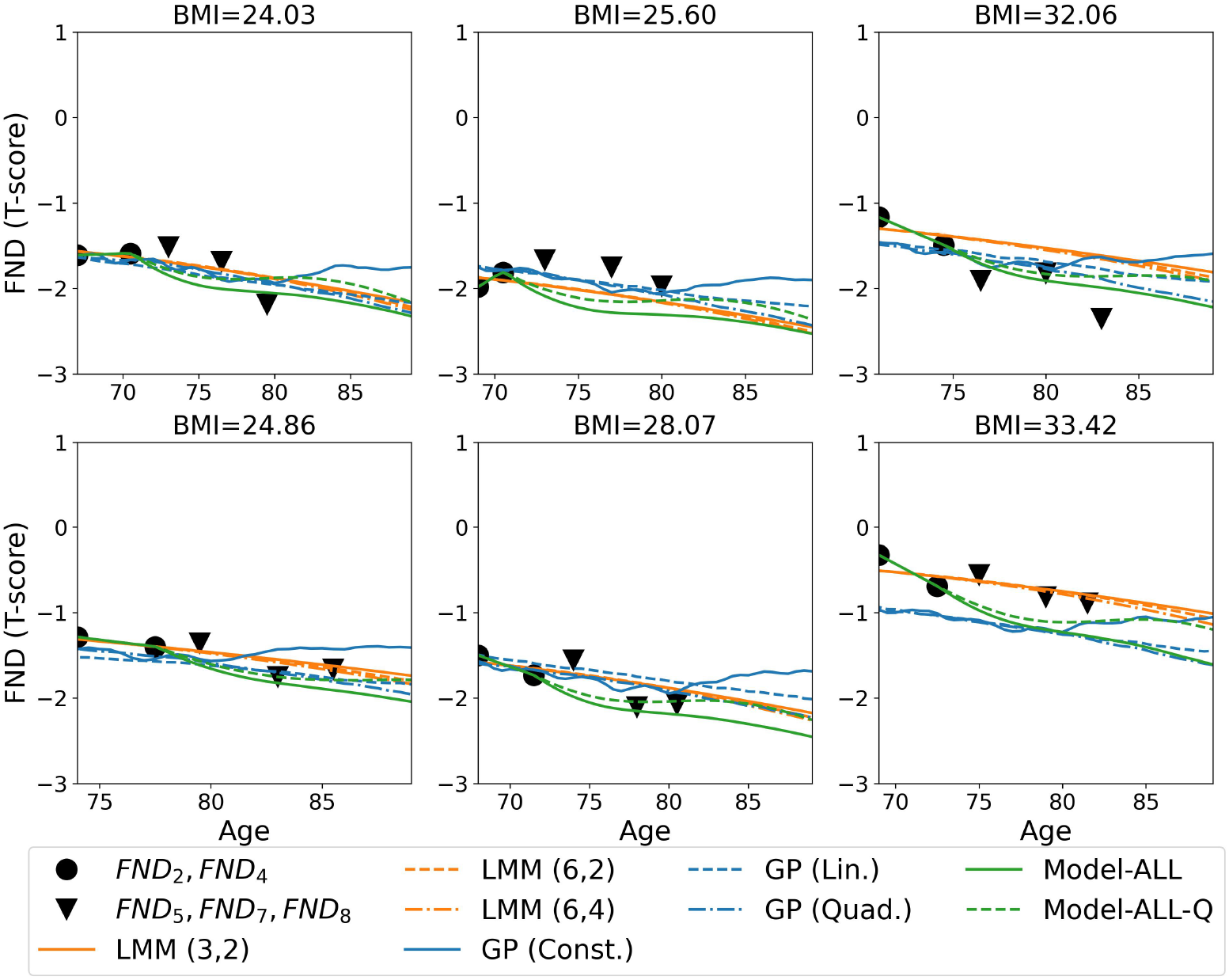
Comparing the mean predicted trajectories with LMMs, GPs, Model-ALL, and Model-ALL-Q, with 𝒥 = {5, 8} for the later two approaches. The six test participants are the same as those considered in Figures 2, 3 and 4. Each plot depicts the phenotype measurements at the initial time-points *t*_2_, *t*_4_ (black dots) which are fed to the models as input, and future time-points *t*_5_, *t*_7_, *t*_8_ (black triangles). The mean trajectories with the LMM models are shown in orange, with the GPs are shown in blue, and the DyViA-GAN are shown in green. The BMI of the participant is listed on the top of each plot.

#### 3.4.3 Gaussian process regression

We consider GP models with mixed effects. For individuals *i* and *j* whose FND are observed at times *t* and *t*^′^, the mixed kernel is taken as *k*((*i, t*), (*j, t*^′^)) = *k*_pop_(*t, t*^′^) + *δ*_*i,j*_*k*_indiv_(*t, t*^′^), where *k*_pop_ models the population-level dynamics, *k*_indiv_ models the individual-level dynamics, and *δ*_*i,j*_ = 1 only if *i* = *j* (i.e., both measurements are for the same individual). In the present work, we take *k*_pop_ to be the Matérn kernel with parameter *ν* = 2.5 and *k*_indiv_ to be a rougher Matérn kernel with *ν* = 0.5. Furthermore, we consider the mean of GP to be either a constant mean, linear, or quadratic. The coefficients of the mean and kernel length-scales are determined by maximizing the log-likelihood on the (unstratified) training data. Like the LMMs and DyViA-GAN, the predictions with trained GP models are conditioned on the measurements at the first two time-points. We use the nomenclature GP Const., GP Lin., and GP Quad., based on how the mean is modeled.

We observe from the RMSE comparison in Table 6 as well as the predicted mean trajectories in Figure 5 that the GP models perform poorly when compared to LMMs and DyViA-GAN. Unlike the LMMs, the predictions with the GPs are not linear, and appear to possess similar expressivity to DyViA-GAN. Thus, it may be possible to fine-tune the GP, or choose a more sophisticated GP variant to improve its performance. This will be explored in future extensions of this work.

## 4 Discussion

While noting that the concept of *healthspan* is relatively new in geroscience research [5] and there is no universal agreement yet on its definition, the author of a perspective paper in *GeroScience* made two key observations: (ii) “health itself may be better considered as a continuous variable that changes in a dynamic way throughout life”; and (2) “[t]he health trajectory will be different in different individuals, but will generally trend downward with age”.[28] However, understanding and prolonging individual-specific healthspan is challenging for many reasons, including not being timely aware of the trajectories of the different aging phenotypes. Advance knowledge of progression of such an outcome as osteopenia, the precursor of osteoporosis, is crucial, as it can lead to an accelerated reduction in one’s bone healthspan, and DyViA-GAN provides precisely that capability.

While there are several powerful approaches that one could use for the trajectory prediction problem, we used a GAN-based approach motivated by the following considerations:

- GANs allow for the generation of many possible outputs associated with a given input. In other words, it is a probabilistic method versus a deterministic one.
- Deep learning frameworks, such as GANs, allow the easy integration of different data modalities. For example, we can easily handle input comprising CT scans (i.e., images type data) and femoral neck density at different time points (i.e., longitudinal scalar data). Furthermore, we have the flexibility of having a different data type for the predictions. This level of flexibility is seldom granted by more traditional algorithms.
- Once trained, inference/predictions on new data is instantaneous. This is in fact an advantage over more sophisticated generative algorithms such as diffusion models [7].

The present study has, in fact, some distinct advantages. Using only very few initial observations as input, it demonstrated the capability of cGAN-type models to predict the personalized trajectory of a single phenotype **–** femoral neck BMD **–** that is particularly relevant to one’s healthspan, and increasingly so as one continues to age. The generators of the proposed DyViA-GAN takes as input the (scaled) age variable *t* and the output generated is a *continuous* trajectory for the entire age-span, and not just for some discrete time points at which data is available during training/testing. The continuous representation also allows for easy calculation of the gradient (and higher derivatives) of each trajectory using automatic differentiation on neural nets.

Notably, there are several key findings from the results presented in section 3. First, the score-based metric indicates that the 𝒥 = {5} DyViA-GAN models generate more spread-out trajectories. However, this might lead to the generation of outlier trajectories which can skewed statistical predictions. This was observed when using the RMSE metric which clearly showed that 𝒥 = {5, 8} DyViA-GAN models lead to better predictions (in the mean), even when compared with a standard spline-based regression model. Second, the inclusion of additional risk factors, such as BMI, may lead to better personalized predictions in certain situations. Third, the 𝒥 = {5} models lead to generation of simplistic trajectories as compared 𝒥 = {5, 8}, indicating the latter are more suitable to capture nonlinear dynamics. Fourth, filtering the predicted trajectories provides a useful strategy to eliminate outliers, leading to improved predictions for the unseen data from Visit 7. Fifth, the Model-ALL-Q variant of DyViA-GAN is at par with standard LMMs in terms of the RMSE metric while being more expressive when predicting individual-level non-monotone trajectories. The Model-ALL-Q and LMMs outperform the GP models.

According to the U.S. Census Bureau’s 2017 National Population Projections, the year 2030 will mark an important demographic turning point in U.S. history when one in five Americans will be 65 years or older. Clearly, planning for and achieving individual healthspan goals will assume increasing importance in the society. Towards this, a variety of new sources of rich, high-resolution data such as genomics, epigenetics, wearable monitoring devices, etc., is making healthcare more preventive and personalized. Such data also allows for complex interplays among the factors of healthy aging **–** ranging from static to dynamic, and both internal and external **–** to inform the models for predicting a wider and more insightful set of outcomes. For instance, a better understanding of the non-linear relationship between one’s BMI and BMD, over the course of aging, could emerge [11]. Towards this end, the translational capacity of AI-based approaches to preventive and precision medicine including the present platform can be refined and tested in clinical trials.

We understand that there are certain limitations of our study. As future work, our approach could be extended to predict simultaneously the trajectories of multiple phenotypes, say, BMD values of multiple regions of the body or fusion thereof. Moreover, key insights are available from other markers of osteoporosis beyond BMI, which we chose for illustrative purposes and for its easy availability and relevance, e.g., [2]. In fact, DyViA-GAN has the flexibility to include, in addition to static baseline risk factors, also dynamic covariates such as a participant’s physical activity level, chronic inflammation, bone biomarkers, epigenetic changes, etc. The effect of including such covariates in the trajectory prediction problem will be considered in future extensions. We also plan to train the models with longitudinal BMD data from heterogeneous populations, e.g., [22]. The presently generated trajectories are not dynamically updated, which could be revised in the future to allow for intermediate events, lifestyle alterations, and medical interventions. Further, we plan to extend the application to more sophisticated deep learning architectures to build the cGAN generator that accounts for the temporal nature of data (e.g., RNN, LSTM), alternative deep generative frameworks (e.g., [3, 13]), as well as other relevant longitudinal datasets on aging phenotypes and groups to train our generative models (e.g., [12]).

## Acknowledgments

The authors thank the SOF Research Group for the publicly available data.

## Appendix A Additional results

As a companion to the test samples considered in Figure 2, Figure 3, Figure 4, we plot a few (filtered) sample trajectories predicted by DyViA-GAN for each of these test samples. These are shown in Figure A.1, Figure A.2 and Figure A.3 below.

**Figure A.1:**
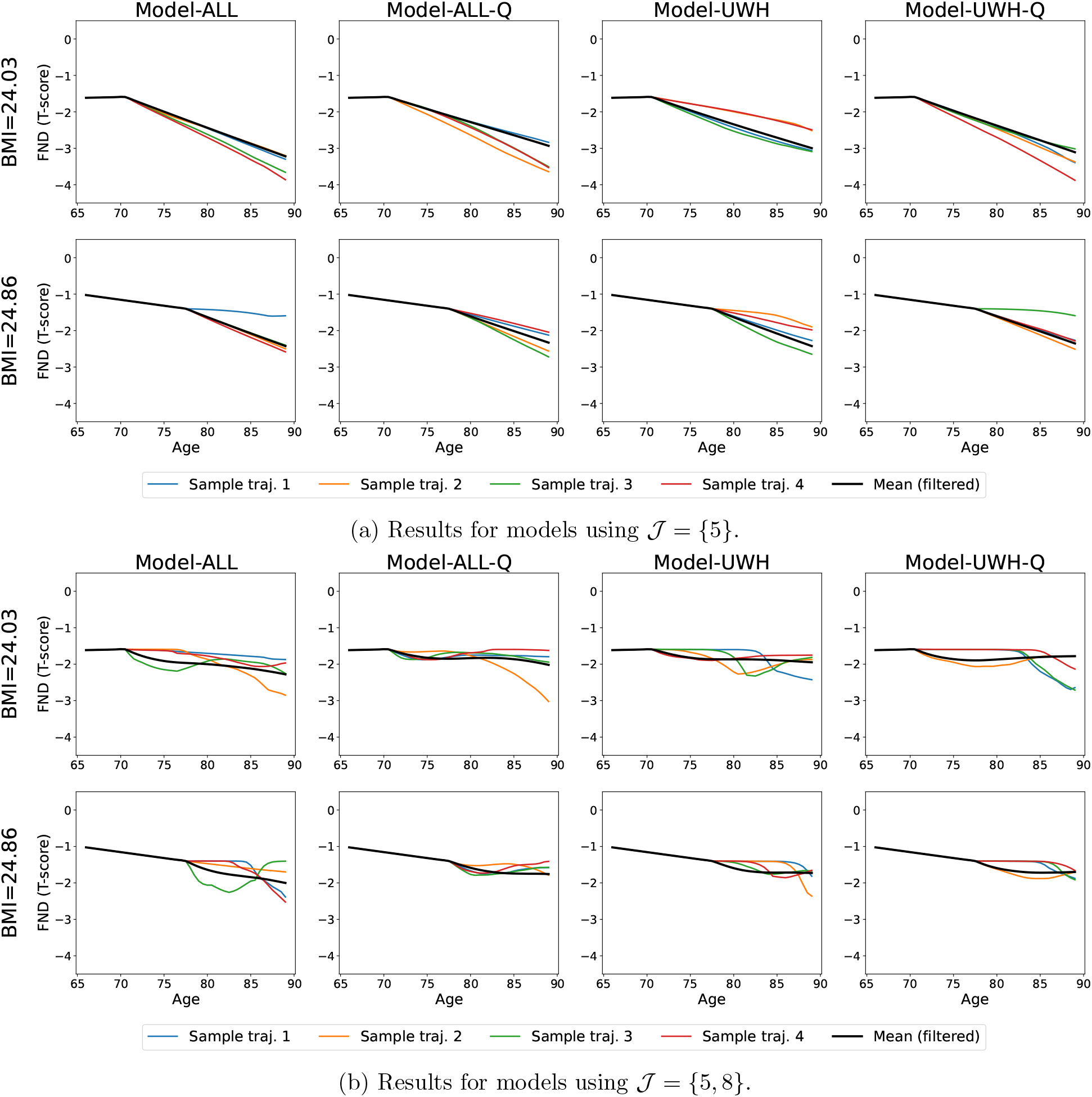
Predictions with models for two test participants from strata UWH. Each plot depicts the (filtered) mean personalized phenotype trajectory (black curve) and four (filtered) sample trajectories. The BMI of the participant is listed on the left of each row.

**Figure A.2:**
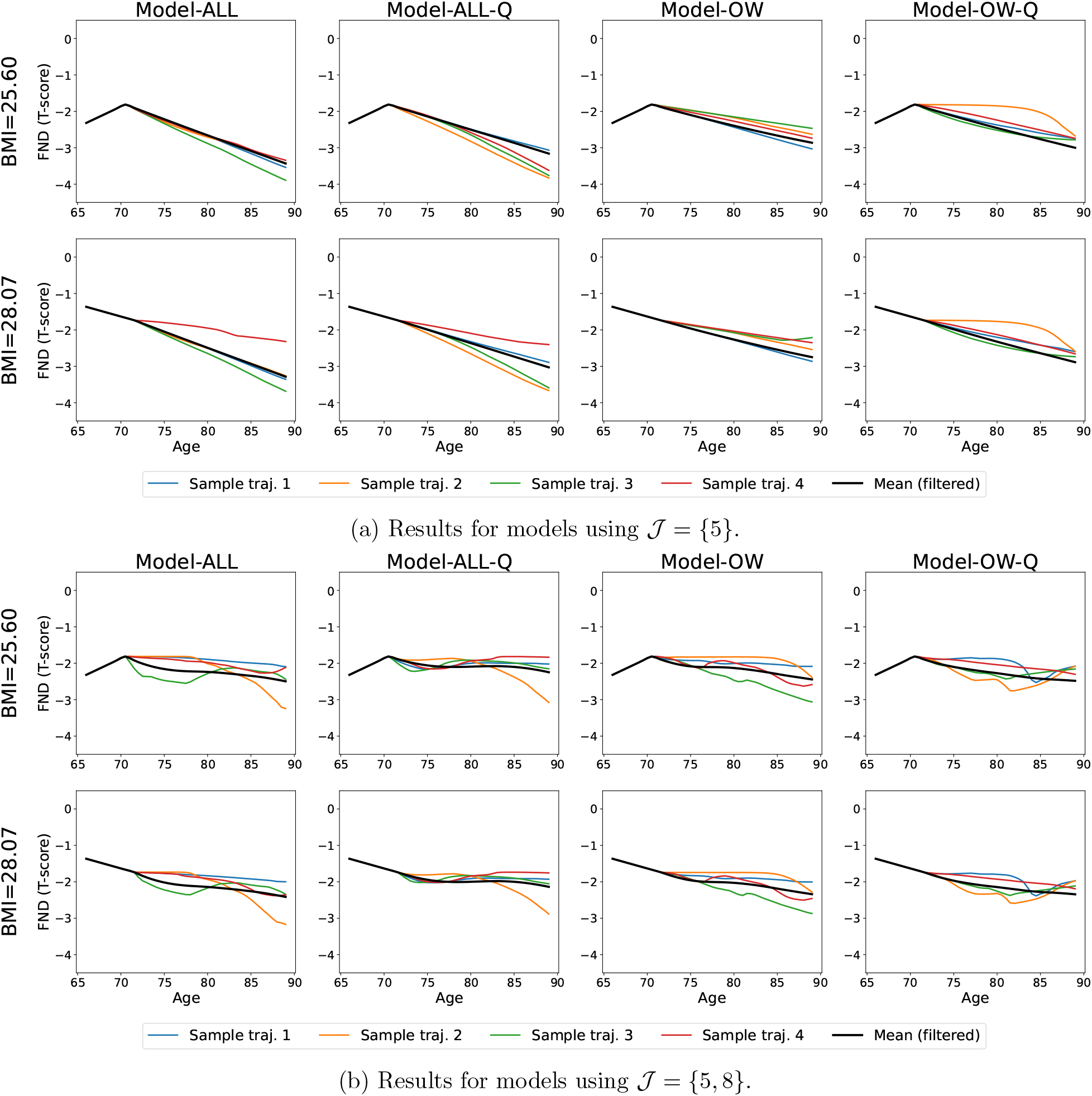
Predictions with models for two test participants from strata OW. Each plot depicts the (filtered) mean personalized phenotype trajectory (black curve) and four (filtered) sample trajectories. The BMI of the participant is listed on the left of each row.

**Figure A.3:**
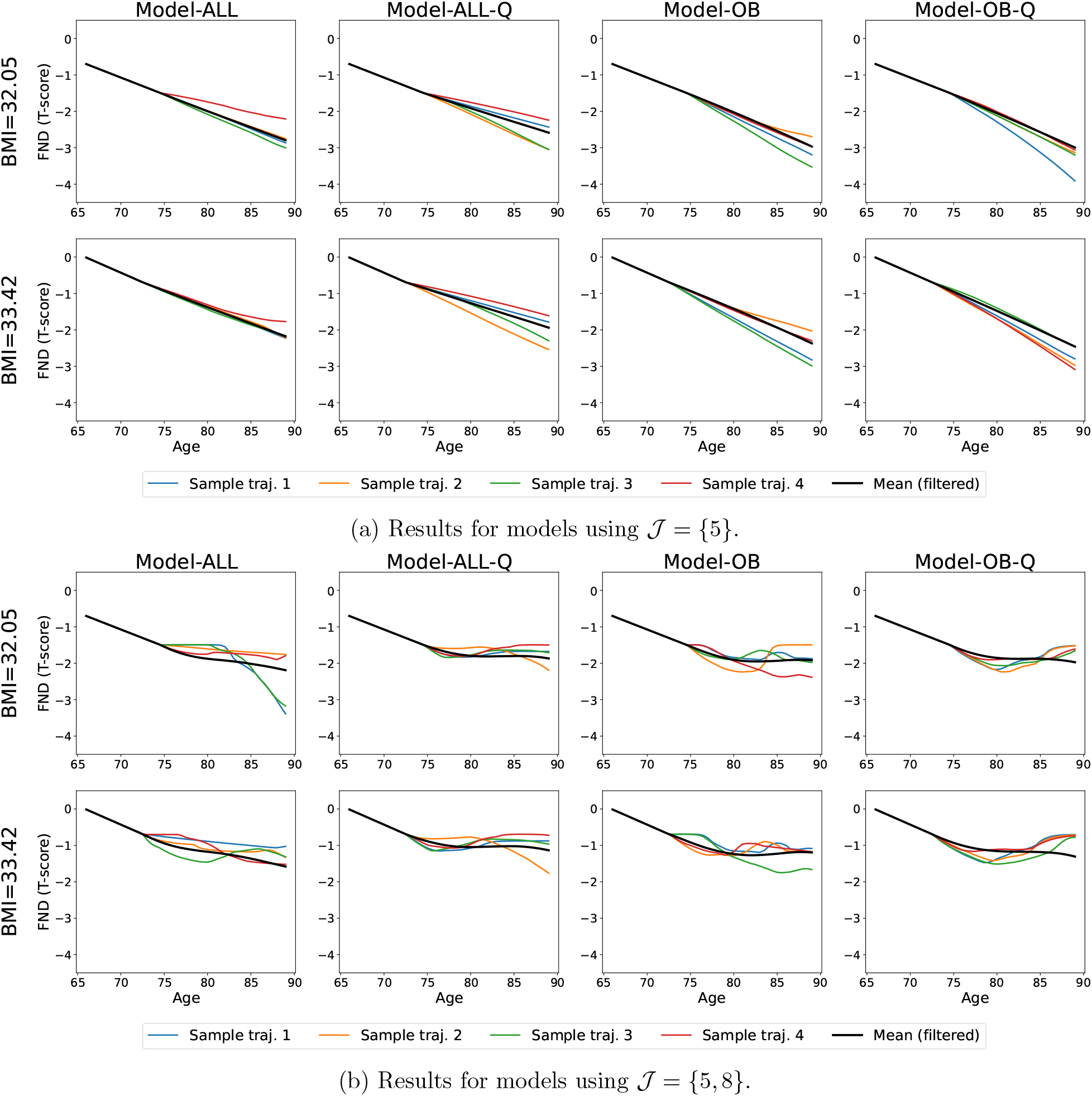
Predictions with models for two test participants from strata OB. Each plot depicts the (filtered) mean personalized phenotype trajectory (black curve) and four (filtered) sample trajectories. The BMI of the participant is listed on the left of each row.

**Figure B.1:**
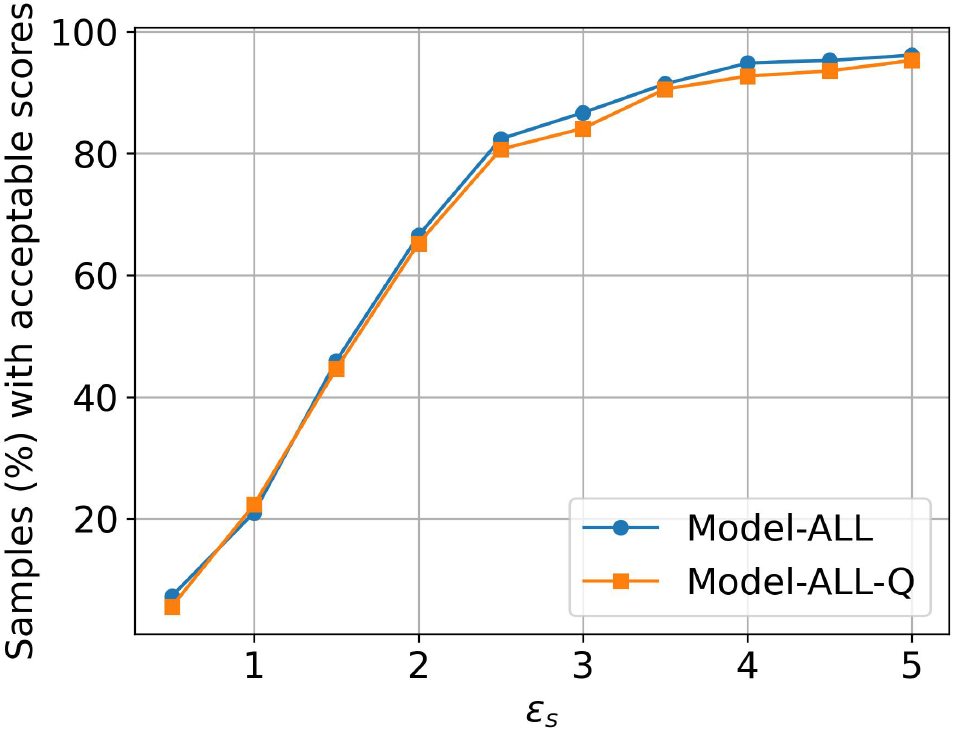
Ablation study to understand the effect of *ϵ*_*s*_ in selecting samples with an acceptable score. The results are shown for Model-ALL (blue curve) and Model-ALL-Q (orange curve), with the y-axis indicating the percentage of test samples flagged as have a suitable score, i.e., 𝒮_*M*_ ≤ *ϵ*_*s*_.

## Appendix B Ablation study: choosing *ϵ*_*s*_ and *ϵ*_*f*_

To determine a suitable value of *ϵ*_*s*_ which is used to define the acceptable scores, we study how the percentage of test samples with an acceptable score, i.e., 𝒮_*M*_ ≤ *ϵ*_*s*_, changes as *ϵ*_*s*_ is increased. Figure Figure B.1 plots this change with Model-ALL and Model-ALL-Q (with 𝒥 = {5, 8}), both of which are trained on the same unstratified dataset. We observe a sharp transition near *ϵ*_*s*_ = 2.5 at which point about 80% of the test sample predictions have acceptable scores. Beyond this threshold, there is a very gradual change in the influence of *ϵ*_*s*_ over the acceptable scores of test samples. Thus, we choose *ϵ*_*s*_ = 2.5 in our experiments.

Next, we determine a suitable value for *ϵ*_*f*_ which is used to filter the samples trajectories on individuals based on DTW distance. In Figure Figure B.2, we plot the fraction of trajectories predicted (by Model-ALL-Q) that are retained for various values of *ϵ*_*f*_. The six colored solid dotted lines show the fractions for the same six test participants considered in the previous section, while the black dashed dotted line shows the median fractions for each value of *ϵ*_*f*_ considered. We observe a sharp transition near *ϵ*_*f*_ = 2.0, at which roughly 88% of trajectories are retained. The fraction increase more gradually beyond this value of *ϵ*_*f*_. This motivates our choice for *ϵ*_*f*_ = 2.0 in our experiments.

**Figure B.2:**
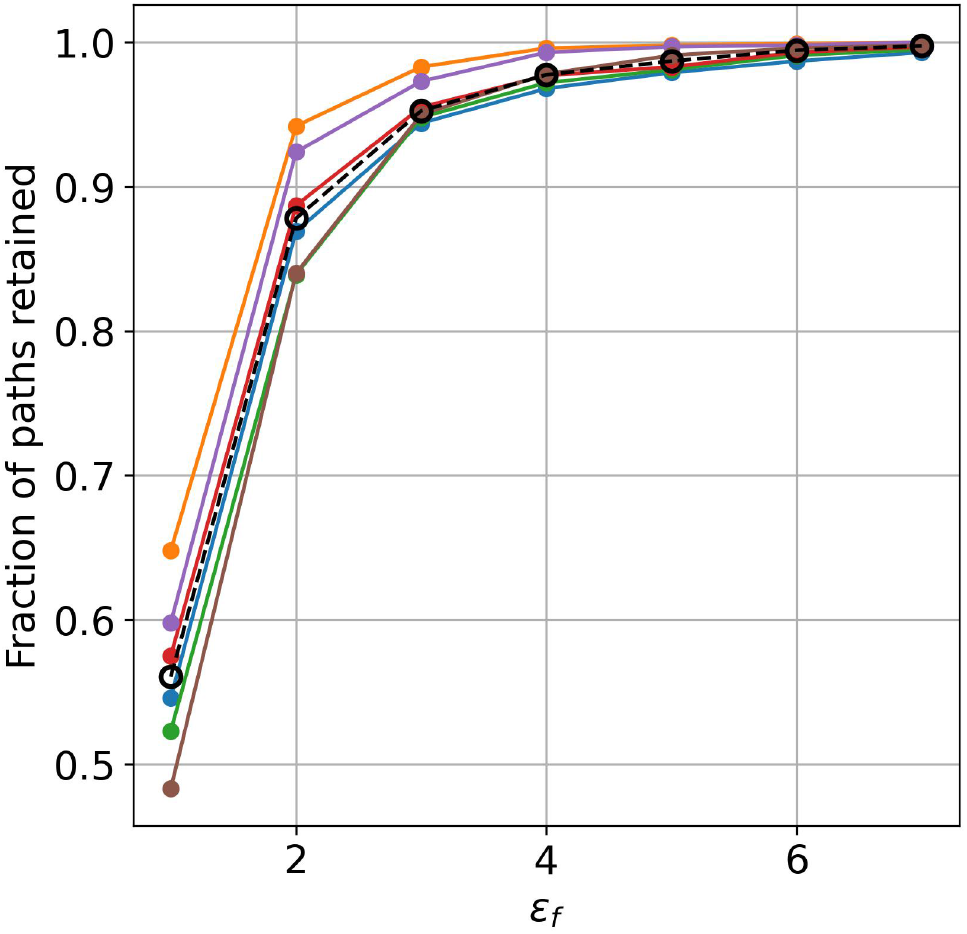
Ablation study to understand the effect of *ϵ*_*f*_ in filtering trajectories based on the DTW distance for different individuals. The colored solid dotted lines show the fraction of predicted trajectories retained as *ϵ*_*f*_ is varied for six test participants for illustrative purposes. The black dashed dotted line shows the median fractions for each value of *ϵ*_*f*_ considered.

## Footnotes

* https://www.nia.nih.gov/health/osteoporosis/osteoporosis

